# *notum1*, acting downstream of pitx2, is essential for proper eye and craniofacial development

**DOI:** 10.1101/687798

**Authors:** Kathryn E. Hendee, Elena A. Sorokina, Sanaa S. Muheisen, Ross F. Collery, Elena V. Semina

## Abstract

Axenfeld-Rieger syndrome (ARS) is a rare autosomal dominant developmental disorder characterized by ocular anterior chamber anomalies with an increased risk of glaucoma and systemic defects. Mutations in the transcription factor *PITX2* were the first identified genetic cause of ARS. Despite the developmental importance of *PITX2* and its role in ARS, the pathways downstream of PITX2 have yet to be fully characterized. Comparative transcriptome analyses involving *pitx2*-enriched cell populations isolated via fluorescence activated cell sorting of tissues expressing (*Tg(-2.6pitx2-CE4:GFP))* reporter in wild-type and *pitx2*^*M64\**^ mutant zebrafish embryos identified the highly down-regulated target *notum1b*, an ortholog of human *NOTUM* encoding a secreted carboxylesterase that cleaves a necessary palmitoleate moiety from WNT proteins. Further experiments confirmed a decrease in *notum1b* and identified down-regulation of another NOTUM ortholog, *notum1a*, in the developing mutant eye. CRISPR-generated permanent double knockout zebrafish lines of *notum1b* and *notum1a, notum1−/−,* displayed defects in craniofacial and ocular development, including corneal defects, small lenses, increased sizes of the anterior and posterior chambers, and anomalies in teeth development. Analysis of head transcriptome of *notum1−/−* zebrafish in comparison to wild-type predicted an up-regulation of the WNT pathway. We present *NOTUM/notum1* as an important factor in ocular and craniofacial development and a novel downstream member of the PITX2/pitx2 pathway.

## INTRODUCTION

Paired-like homeodomain 2 (*PITX2* [MIM 601542]) is a *bicoid*-like homeodomain transcription factor that plays a key role in early development (Semina et al., 1996). *PITX2* is the first causative gene associated with Axenfeld-Rieger syndrome, type 1 (ARS [MIM #180500]), a rare developmental disorder characterized by particular ocular anomalies and systemic defects (Reis & Semina, 2011; Reis et al., 2012; Semina et al., 1996). Ocular features present as any combination of iris processes, corectopia/polycoria, iris hypoplasia, and posterior embryotoxon, along with a 50% risk of developing glaucoma (Alward, 2000; Reis & Semina, 2011). Classical accompanying systemic defects include maxillary hypoplasia, micro- or hypodontia, peg-like incisors, and redundant periumbilical skin (Reis & Semina, 2011; Reis et al., 2012; Strungaru, Dinu, & Walter, 2007; Tumer & Bach-Holm, 2009). Complete gene deletion alleles support *PITX2* haploinsufficiency as a mechanism of ARS (Flomen et al., 1998; Reis et al., 2012; Semina et al., 1996; Strungaru et al., 2007; Tumer & Bach-Holm, 2009). A majority of *PITX2* mutations affect the homeodomain and C-terminal regions shared by the three main mammalian transcripts *PITX2-A, -B*, and *-C* through deletions, splicing, or missense alleles, thus resulting in partial or complete loss of function due to the disruption of all PITX2 isoforms (Cox et al., 2002; Reis et al., 2012; Strungaru et al., 2007; Tumer & Bach-Holm, 2009).

PITX2 function is highly conserved during vertebrate development. In mice, *Pitx2* is expressed in tissues affected by ARS, namely the periocular mesenchyme, maxillary and mandibular epithelia, dental lamina, and umbilicus (Semina et al., 1996). Homozygosity for *Pitx2* null or hypomorphic alleles is embryonic lethal, and among the multiple severe anomalies observed are ARS-related features including irregular mandibular and maxillary facial prominences, arrested tooth development, absent ocular muscles, and abnormal eye placement and anterior segment structures (Gage, Suh, & Camper, 1999; Kitamura et al., 1999; Lin et al., 1999; Lu, Pressman, Dyer, Johnson, & Martin, 1999). In addition, heterozygous *Pitx2* mutant mice were recently demonstrated to manifest the ocular features of ARS (Chen & Gage, 2016). Vertebrate conservation also extends to zebrafish where two isoforms, *pitx2a* and *pitx2c*, have shown to be positionally and structurally conserved with human *PITX2A* and *PITX2C* respectively (Volkmann et al., 2011). We recently generated and characterized a *pitx2*-deficient zebrafish line. Observed ocular defects in homozygous *pitx2* mutants included an underdeveloped anterior chamber with abnormalities in the cornea, iris and iridocorneal angle structures. Additionally, the pharyngeal arch cartilages of the lower jaw exhibited structural and positional defects. These features emulated several key features observed in ARS as well as rare phenotypes connected with *PITX2,* namely ring dermoid of the cornea (Hendee et al., 2018).

Despite the developmental importance of *PITX2* and its role in ARS, the pathways downstream of PITX2 have yet to be fully characterized. As a transcription factor, Pitx2 is typically considered an activator of expression of its target genes (Amendt, Sutherland, & Russo, 1999). Among identified Pitx2 targets, a subset most relevant to the developmental eye and craniofacial phenotypes includes Dickkopf 2 (*Dkk2*), an extracellular antagonist of canonical Wnt-β-catenin signaling that was the first identified downstream target of Pitx2 in periocular mesenchyme during anterior segment, particularly cornea, development (Gage, Qian, Wu, & Rosenberg, 2008). Pitx2 was also determined to be required for normal *Tfap2b* expression within the neural crest, which is necessary for corneal endothelial development and the establishment of corneal angiogenic privilege (Chen et al., 2016). In developing craniofacial structures, distal-less homeobox 2 (*Dlx2*) was shown to be downstream of Pitx2 within the dental epithelium during tooth development (Green et al., 2001). Transcriptome analysis of ocular tissues from our *pitx2*^*M64\**^ zebrafish line suggested several genes, namely collagens (e.g., I and V) and Wnts (e.g., 3 and 10a), as additional potential downstream targets of pitx2 (Hendee et al., 2018). Further dissection of the PITX2/pitx2 pathway through identification of its critical downstream targets is needed to better understand its role in embryonic development and human disease.

In this manuscript we describe the comparative transcriptome analyses involving *pitx2*-enriched cell populations from wild-type and *pitx2*^*M64\**^ zebrafish embryos and the identification of the highly down-regulated target *notum1b,* an ortholog of human *NOTUM* encoding a secreted carboxylesterase that cleaves a necessary palmitoleate moiety from WNT proteins (Kakugawa et al., 2015; Zhang et al., 2015). We demonstrate that *notum1 (notum1a−/−; notum1b−/−)* deficiency in zebrafish results in ocular and craniofacial anomalies overlapping those of *pitx2* deficiency. Thus, we present *NOTUM/notum1* as an important factor in ocular and craniofacial development and a novel downstream member of the PITX2/pitx2 pathway.

## MATERIALS AND METHODS

### Transcriptome analysis in pitx2^M64*^ and notum1-deficient embryos

The mutant *pitx2* allele c.190_197del8 (p.(M64*)) was previously described (Hendee et al., 2018). This variant was produced on a zebrafish reporter line where GFP is controlled by conserved upstream regulatory element 4 of *pitx2* (*Tg(-2.6pitx2-CE4:GFP))* (Volkmann et al., 2011). Head tissues were micro-dissected from a mix of (*CE4:GFP):pitx2*^*M64\**^ heterozygous (50%) and homozygous (50%) embryos along with (*CE4:GFP):WT* control embryos at 24-hours post fertilization (hpf). Heads were dissociated into a single cell suspension in trypsin, spun down, washed once with Ringer’s solution, then resuspended in 2 mL of Ringer’s plus 5% fetal bovine serum (FBS) and 1 ul/mL propidium iodide. GFP-positive and negative cell populations were isolated using fluorescence-activated cell sorting (FACS) on a FACSAriaII special order system Cell Sorter (BD Biosciences, San Jose, CA). RNA was extracted from FACS-collected cells with TRIzol and the Direct-zol RNA MiniPrep Kit (Zymo Research, Irvine, CA). RNA quality was assessed on an Agilent 2100 Bioanalyzer utilizing the Agilent RNA 6000 Pico Kit (Agilent Technologies, Santa Clara, CA) as well as a NanoDrop 2000 UV-Vis Spectrophotometer (Thermo Fisher Scientific, Waltham, MA). For wild-type GFP-positive and negative cell populations, six independent collections generated four RNA samples for each condition, with RNA Integrity Numbers (RINs) of 9.2, 8.3, 7.4, and 7.8 for GFP-positive and 8.9, 8, 9.2, and 9.2 for GPF-negative cells. Likewise, RNA samples with RINs 8.8, 9.1, 9.5, and 8.5 for GFP-positive and 8.1, 9.7, 8.5, and 8.2 for GFP-negative cells were obtained from six independent (*CE4:GFP):pitx2*^*M64\**^ collections. RNA was submitted to OakLabs (Hennigsdorf, Germany) for transcriptome analysis using the zebrafish-specific microarray platform ArrayXS Zebrafish v1 Zv9 with 60,023 target IDs. OakLabs also performed statistical analysis, including quantile normalization with ranked median quantiles, conversion to log(2) values to calculate control versus mutant means and standard deviations per target, and significance testing using Welch’s t-test. Differentially expressed targets were defined as having a difference in mean log(2) values > +1 or < −1 and a p-value < 0.05. Zebrafish-specific transcripts were annotated with human orthologs as previously described (Hendee et al., 2018). The list of human orthologs was analyzed for pathway enrichment by Ingenuity Pathway Analysis software (Qiagen, Hilden, Germany).

*notum1a, notum1b,* and *dkk2* transcript levels were verified by real-time quantitative reverse transcription PCR using transcript-specific primers (Table S1), SYBR Green PCR Master Mix (Applied Biosystems, Waltham, MA) and CFX Connect and CFX96 Touch Real-Time PCR Detection Systems (Bio Rad, Hercules, CA). Equal concentrations of RNA from 24-hpf (*CE4:GFP):WT* and (*CE4:GFP):pitx2^M64*^* GFP-positive cells, GPF-negative cells, and whole eyes that did not undergo FACS were used to generate cDNA with SuperScript III reverse transcriptase (Thermo Fisher). Each qPCR experiment included the following conditions at 150 pg cDNA per replicate: 24-hpf (*CE4:GFP):WT* GFP-positive, GPF-negative, and whole eye cDNA; 24-hpf (*CE4:GFP):pitx2*^*M64\**^ GFP-positive, GPF-negative, and whole eye cDNA; and no template control which excluded primers. All samples were normalized to β-actin and run in triplicate to obtain average threshold cycle (*CT*) values. Total fold changes and standard deviations were calculated as the average of three independent biological repeats via the 2^−ΔΔ*CT*^ model.

A second microarray was performed on wild-type and *notum1*-deficient (*notum1a−/−; notum1b−/−*) whole heads collected at 24-hpf. Independent collections generated three RNA samples for wild-types and four RNA samples for *notum1* mutants, which were extracted and assessed as described above. RNA samples with RINs of 8.7, 9.6, and 9.1 for wild-type and 9.5, 9.5, 9.5, and 7.9 for *notum1*-deficient heads were submitted to OakLabs for the same transcriptome and statistical analysis, and the resulting data was annotated and examined in the same manner with the addition of cross-referencing against the first *pitx2*^*M64\**^ array.

### Development of notum1a−/−;notum1b−/− mutant lines via CRISPR/Cas9 genome editing

Zebrafish maintenance and staging were performed as previously described (Liu & Semina, 2012). CRISPR/Cas9 genome editing (Hwang et al., 2013) was utilized to target the regions encoding the catalytic domains of *notum1a* and *notum1b*. Single guide RNA (sgRNA) oligonucleotides were designed using ZiFiT Targeter software (http://zifit.partners.org/ZiFiT/ChoiceMenu.aspx) (Sander et al., 2010; Sander, Zaback, Joung, Voytas, & Dobbs, 2007) and the Broad Institute sgRNA Designer (https://www.genscript.com/gRNA-design-tool.html) (Table S1). MLM3613 (Addgene plasmid # 42251, Addgene, Cambridge, MA) and DR274 (Addgene plasmid #42250) plasmids (gifts from Keith Joung) were utilized for Cas9 mRNA and sgRNA synthesis (Hwang et al., 2013; Sander et al., 2010). Cas9 mRNA, sgRNAs, and nanos for visualization of germline incorporation (Dong, Dong, Jia, Cao, & Zhao, 2014; Koprunner, Thisse, Thisse, & Raz, 2001) were injected into 1-4 cell stage wild-type embryos at concentrations of 27.2 ng/μl, 2.7 ng/μl, and 54.4 ng/μl respectively, generating mosaic founders that produced single and double F1 heterozygous *notum1a* and *notum1b* mutant embryos. PCR amplification and sequencing of F1 offspring with *notum1a*-specific primers (Table S1) resulted in five frameshift mutations: c.641delA, p.(Lys214Argfs*4); c.640_641delAA, p.(Lys214Glyfs*13); c.640_644delAAGGA, p.(Lys214Glyfs*12); c.743_749delinsTGCAAGACCTTTTGGAA, p.(Asp248Valfs*39); and c.743_746delinsTGTCTT, p.(Asp248Valfs*104). Two *notum1a* frameshift alleles also had additional mutations in cis but predicted to be downstream of protein truncation: c.740_745delTGGACT, p.(Val247_Ser249delinsAla) was in cis with c.640_644delAAGGA, p.(Lys214Glyfs*12); and c.743_748delACTCGG, p.(Asp248_Ser249del) was in cis with c.640_641delAA, p.(Lys214Glyfs*13). The same analysis with *notum1b*-specific primers (Table S1) identified three frameshift mutations— c.556_557insTGATCTCTGATCATT, p.(Gln186Leufs*6); c.557_572delinsTCTTAACT, p.(Gln186Leufs*7); c.558_565delGGAGGTCA, p.(Gln186Hisfs*7)— and one in-frame deletion— c.555_566delTCAGGAGGTCAT, p.(Gln186_Ile189del). *notum1a* lines used for analysis were compound heterozygous for the c.743_749delinsTGCAAGACCTTTTGGAA, p.(Asp248Valfs*39) and c.640_641delAA, p.(Lys214Glyfs*13) alleles (*notum1a^*D248Vfs*39*^/*notum1a*^*K214Gfs\**^*^13^). *notum1b* lines were homozygous for the c.558_565delGGAGGTCA, p.(Gln186Hisfs*7) allele (*notum1b^Q186Hfs*7^*). *notum1* lines were homozygous for both the *notum1a* allele c.640_644delAAGGA, p.(Lys214Glyfs*12) and the *notum1b* allele c.557_572delinsTCTTAACT, p.(Gln186Leufs*7) (*notum1a^K214Gfs*12^; notum1bQ186Lfs*7*).

### Characterization of embryonic and adult phenotypes in notum1 mutants

Alcian Blue staining was performed in both acidic and acid-free conditions. Acidic Alcian Blue followed the protocol of Barrallo-Gimeno et al. as previously described (Barrallo-Gimeno, Holzschuh, Driever, & Knapik, 2004; Hendee et al., 2018). Acid-free Alcian Blue and Alizarin Red dual staining, for cartilage and bone respectively, was conducted as per Walker and Kimmel (Walker & Kimmel, 2007) with minor deviations. A stock of 0.4% Alcian Blue was prepared by dissolving Alcian Blue 8GX (Sigma-Aldrich, St. Louis, MO) in 50% ethanol (EtOH) at 37°C then adding 100% EtOH to reach a final concentration of 70% EtOH. 100 ml of Part A of the dual stain [0.02% Alcian Blue] was made by combining 60 mM magnesium chloride, 100% EtOH, and 0.4% Alcian Blue stock. The stock of Part B, 0.5% Alizarin Red, was prepared by dissolving Alizarin Red S (Sigma-Aldrich) in water. 5-dpf embryos were collected and fixed in 4% paraformaldehyde in 1x diethyl pyrocarbonate phosphate-buffered saline (DEPC-PBS) at room temperature rocking for 2 hrs, overnight at 4°C, and 1 hr at room temperature before dehydration with 50% EtOH on a rocker for 10 mins. 1 ml of Part A and 10 μl of Part B were combined per tube of embryos just prior to staining, and embryos were rocked in the dual stain solution overnight. Embryos were rinsed with water, bleached in equal parts 3% hydrogen peroxide (H_2_O_2_) and 2% potassium hydroxide (KOH) for 20 mins with open tube lids, and cleared with a 2.5-hr rocking wash in 20% glycerol and 0.25% KOH followed by an overnight wash in 50% glycerol and 0.25% KOH. Embryos were imaged and stored at 4°C in 50% glycerol and 0.1% KOH. Flat-mount dissection techniques were provided courtesy of Xu et al. (Xu et al., 2018). Craniofacial structures were identified as presented by Piotrowski et al. and Xu et al. (Piotrowski et al., 1996; Xu et al., 2018). Whole-mount and flat-mount images were obtained on a Nikon SMZ1500 dissection microscope (Nikon Instruments Inc., Melville, NY).

For histology, several ages of fish were preserved in 10% formalin or Davidson’s solution [glacial acetic acid, 95% ethyl alcohol, 10% neutral buffered formalin, distilled water] and submitted to the Children’s Research Institute Histology Core at the Medical College of Wisconsin for paraffin sectioning: 5-, 8-, 10-, and 14-dpf (for wild-type numbers were 5, 5, 7, and 4 fish; for *notum1*-deficient, numbers were 5, 6, 6, and 12 fish, respectively); 6.5-, 8.5-, and 13-months post fertilization [mpf] (wild-type: 2, 3, and 2 fish);and 7- and 14-mpf (*notum1*-deficient: 6 and 3 fish). Hematoxylin and eosin (H&E) staining, Periodic acid Schiff (PAS) staining, and immunohistochemistry were performed on fresh slides per standard protocols as previously described (Hendee et al., 2018). H&E and PAS slides were imaged on a NanoZoomer digital slide scanner, and images were viewed with NDP.view2 viewing software (Hamamatsu, Hamamatsu City, Japan). Immunohistochemistry was performed at most collected stages (8-, 10-, and 14-dpf; 6.5-, 7-, 8.5-, 13-, and 14-mpf) with the following antibodies: primary—corneal keratan sulfate proteoglycan (CKS; 1:125; MAB2022; EMD Millipore, Billerica, MA), N-cadherin (cdh2; 1:50; GTX125962; GeneTex, Inc., Irvine, CA), and DAPI (1:1000; 62247; Thermo Fisher); secondary—donkey anti-rabbit Alexa Fluor 488 IgG (1:100; A-21206; Thermo Fisher), and donkey anti-mouse Alexa Fluor 568 IgG (1:100; A-10037; Thermo Fisher). Images were taken on an AxioImager.Z1 microscope with an AxioCam MRc5 camera and ZEN 2 software (Zeiss, Thornwood, NY).

*In situ* hybridization (ISH) and RNAscope were performed as previously described (Hendee et al., 2018; Liu & Semina, 2012). Transcript-specific primers were utilized for contig mapping of *notum1b* as well as, along with DIG RNA labeling mix and T7 RNA polymerase (Roche Diagnostics Corporation, Indianapolis, IN), generation of one *notum1a* and four *notum1b* digoxigenin-labeled anti-sense RNA probes (*notum1a*: 623 bp; *notum1b:* 756, 1424, 763, and 770 bp) (Table S1). Whole mount embryos were imaged as described above. RNAscope probes Dr-notum1a-C2 (499601-C2; Advanced Cell Diagnostics, Newark, CA) and Dr-notum1b-C3 (499671-C3; Advanced Cell Diagnostics) were obtained and used as described by Gross-Thebing et al. (Gross-Thebing, Paksa, & Raz, 2014). Stained embryos were prepared for cryosectioning with two PBS washes, two overnight infiltrations at 4°C, first with 30% sucrose/PBS then Tissue-Tek O.C.T. Compound (Sakura Finetek USA, Inc., Torrance, CA), and embedding and freezing in cryomolds. 10-micron sections were obtained using a Microm HM550 cryostat (Thermo Fisher), melted onto charged glass slides, mounted with ProLong Gold antifade reagent (Invitrogen, Waltham, MA), and imaged as described.

Spectral domain-optical coherence tomography (SD-OCT) was performed as per Collery et al. (Collery, Veth, Dubis, Carroll, & Link, 2014). 7-mpf wild-type and 7 and 8-mpf *notum1*-deficient adults (5, 6, and 2 fish respectively) were anesthetized and their eyes imaged using a Bioptigen Envisu R2200 SD-OCT imaging system with a 12 mm telecentric lens (Bioptigen, Morrisville, NC). Collected metrics include body axis length, axial length, lens diameter, corneal thickness, vitreous chamber depth, anterior chamber area, retinal thickness, and relative refractive error. Relative refractive error was calculated by comparing the location of the predicted focal point of the lens and the actual retinal radius of the eye using the formula 1−(r/(lr × 2.324)), where r is the retinal radius, and lr is the lens radius. Positive values are relatively hyperopic, and negative values are relatively myopic. Images were assembled using InVivoVue software (Leica Microsystems, Buffalo Grove, IL), and ImageJ (Rasband, W.S., ImageJ, U. S. National Institutes of Health, Bethesda, Maryland, USA, https://imagej.nih.gov/ij/,1997-2018). Adjacent frames of volume scans were assembled as movie files to show the spatial distribution of aberrant features of interest in the *notum1−/−* cornea using ImageJ. Eye measurements were processed using Microsoft Excel (Microsoft, Redmond, WA) and graphed using GraphPad Prism (GraphPad, La Jolla, CA).

Electron microscopy was performed as previously described (Hendee et al., 2018; Soules & Link, 2005). 8.5-mpf wild-type and 8-mpf *notum1*-deficient heads were bisected, fixed, and submitted to the Electron Microscopy Facility at the Medical College of Wisconsin for embedding. Transverse, semi-thin (1 μM) sections were cut with a DiATOME Histo diamond knife (HI10213; DiATOME, Hatfield, PA) on a PowerTome XL microtome (Boeckeler Instruments, Tucson, AZ) and collected throughout the anterior chamber until maximum lens diameter was reached. Ultrathin sections (70–80 nm) were taken at the central anterior chamber and cornea with a DiATOME Ultra 45° diamond knife (MT7376; DiATOME), collected on copper hexagonal mesh coated grids (G200H-Cu; Electron Microscopy Sciences, Hatfield, PA) and stained with uranyl acetate and lead citrate for contrast. Images were captured using a Hitachi H600 TEM microscope (Hitachi, Tokyo, Japan).

## RESULTS

### Identification of differentially expressed transcripts in zebrafish pitx2-deficient tissues

To enrich for transcripts co-expressed with *pitx2*, transcriptome analysis was performed on fluorescently sorted cell populations from 24-hpf head tissues micro-dissected from (*CE4:GFP):WT* control and mutant (a mix of 50% heterozygous/50% homozygous) (*CE4:GFP):pitx2*^*M64\**^ embryos. (*CE4:GFP):WT* utilizes one of the *PITX2/pitx2* enhancers (CE4) and was previously described (Volkmann et al., 2011). A comparison between (*CE4:GFP):WT* and (*CE4:GFP):pitx2*^*M64\**^ GFP-positive cell populations identified significantly differentially expressed targets: 131 up-regulated and 96 down-regulated transcripts in the (*CE4:GFP):pitx2*^*M64\**^ mutant versus the wild-type control. Of these, 112 up-regulated and 82 down-regulated transcripts had recognized human orthologs that, when mapped by Ingenuity Pathway Analysis (IPA), indicated inactivity of adrenomedullin and Protein Kinase A (PKA) signaling and enriched activity of the GP6 signaling pathway (Table S2). More broadly, increased activity was predicted in processes including concentration of dopamine, brain growth, and development of the head, sensory organ, and abdomen, whereas decreased activity was noted in pathways such as cellular attachment and kidney development (Table S2). Full microarray data are available from the ArrayExpress database at EMBL-EBI (www.ebi.ac.uk/arrayexpress; Accession number: pending).

Differentially expressed transcripts were sorted by p-value significance and magnitude of fold change to generate a list of top 50 targets (Figure 1A). The top downregulated candidate (subsequently identified in both downregulated adrenomedullin and PKA signaling pathways) was notum pectinacetylesterase homolog 1b (*notum1b*). Transcript levels of *notum1b* were confirmed to be significantly downregulated in the 24-hpf (*CE4:GFP):pitx2*^*M64\**^ GFP-positive population by qRT-PCR (Figure 1B). Similar analysis of a *notum1b* homolog, *notum1a*, did not show significantly reduced transcript level in the (*CE4:GFP):pitx2*^*M64\**^ GFP-positive cells, in agreement with *notum1a* not meeting the significance threshold of expression fold change on the microarray (Figure 1B). *dkk2* also did not display significant differential expression according to the microarray, though there was a trend toward decreased expression in (*CE4:GFP):pitx2*^*M64\**^ GFP-positive cells. qRT-PCR revealed *dkk2* to be significantly down-regulated in this cell population; thus, *notum1b* and *dkk2* transcript levels followed the same pattern (Figure 1B). Interestingly, *notum1b, notum1a,* and *dkk2* were all significantly down-regulated in 24-hpf (*CE4:GFP):pitx2*^*M64\**^ non-sorted whole eyes as compared to wild-type (Figure 1B). Overall, this data suggests that *notum1b* and *notum1a* levels are disrupted in *pitx2*-deficient ocular tissues and present promising candidates for downstream targets of pitx2.

**Figure 1:**
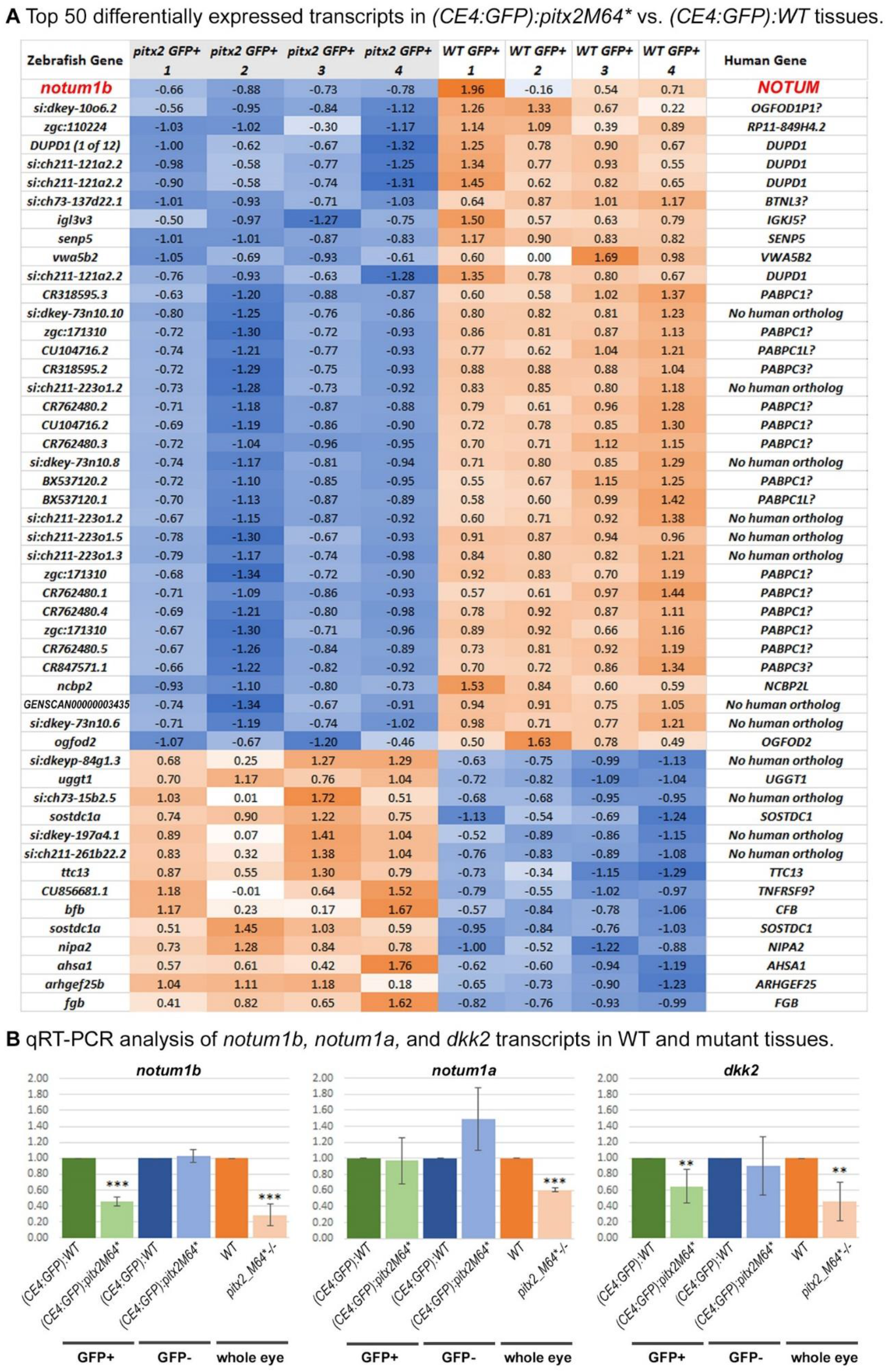
Comparison and qRT-PCR analysis of top differentially expressed transcripts in (*CE4:GFP):pitx2*^*M64\**^ vs. (*CE4:GFP):WT* tissues. **A)** Heatmap of the top 50 significantly down- or up-regulated transcripts in the (*CE4:GFP):pitx2*^*M64\**^ GFP-positive vs. the (*CE4:GFP):WT* GFP-positive cell populations. Human orthologs are indicated for all genes unless no orthologous human sequence was identified, with question marks designating ambiguous entries (low-medium level of identity revealed via BLAST with more studies needed to verify relationships). **B)** qRT-PCR analysis of *notum1b, notum1a,* and *dkk2* transcripts in wild-type and *pitx2^M64*^* mutant tissues. Transcript levels of *notum1b* were confirmed to be significantly down-regulated in the 24-hpf (*CE4:GFP):pitx2*^*M64\**^ GFP-positive population. *notum1a*, a homolog of *notum1b,* did not show significantly reduced transcript level in the (*CE4:GFP):pitx2*^*M64\**^ GFP-positive cells, in agreement with *notum1a* not meeting the significance threshold of differential expression on the microarray. *dkk2* was also revealed to be significantly down-regulated in (*CE4:GFP):pitx2*^*M64\**^ GFP-positive cells. *notum1b, notum1a,* and *dkk2* were all significantly down-regulated in 24-hpf (*CE4:GFP):pitx2*^*M64\**^ non-sorted whole eyes as compared to wild-type. *p<.05 ^**^ p<.01^***^ p<.001.

### Identification and characterization of zebrafish notum 1a and notum1b

Human *NOTUM* spans 11 exons, produces a 2,223 bp transcript, and encodes a 496 amino acid (aa) protein. Human *NOTUM* has three homologs in zebrafish: *notum1a, notum1b*, and *notum2*. *notum1a,* a 2,246 bp transcript encoding a 500 aa protein, was the primary homolog previously studied due to its higher encoded peptide identity (74%) and conserved genomic synteny with *NOTUM* as well as its specific gene expression pattern during the first day of development (Flowers, Topczewska, & Topczewski, 2012b). The available database entries for *notum1b* [NM_001002644.1, NP_001002644.1] predict a 2,128 bp, 49 aa protein that does not contain the conserved pentapeptide active site and catalytic triad, thus rendering the protein inactive. To verify the *notum1b* sequence, we prepared total RNA/cDNA using wild-type embryos and amplified overlapping *notum1b* fragments with specific primers (Table S1). The resultant segments were cloned, sequenced and combined into a contig. The obtained *notum1b* sequence shows 64% (319 of 496 AA) identity with the human protein and contains all functional domains (Figure 2A; Figure S1). The sequence is submitted into GenBank (https://www.ncbi.nlm.nih.gov/genbank; Accession number: pending). The third homolog, *notum2,* shares conserved genomic synteny in all fish but is absent from mammals (Cantu, Flowers, & Topczewski, 2013; Flowers, Topczewska, & Topczewski, 2012a). Despite conservation of active sites, *notum2* was not pursued further in these studies due to its lower encoded peptide identity (51%) with *NOTUM* and possible functional divergence (Figure S1) (Cantu et al., 2013).

**Figure 2:**
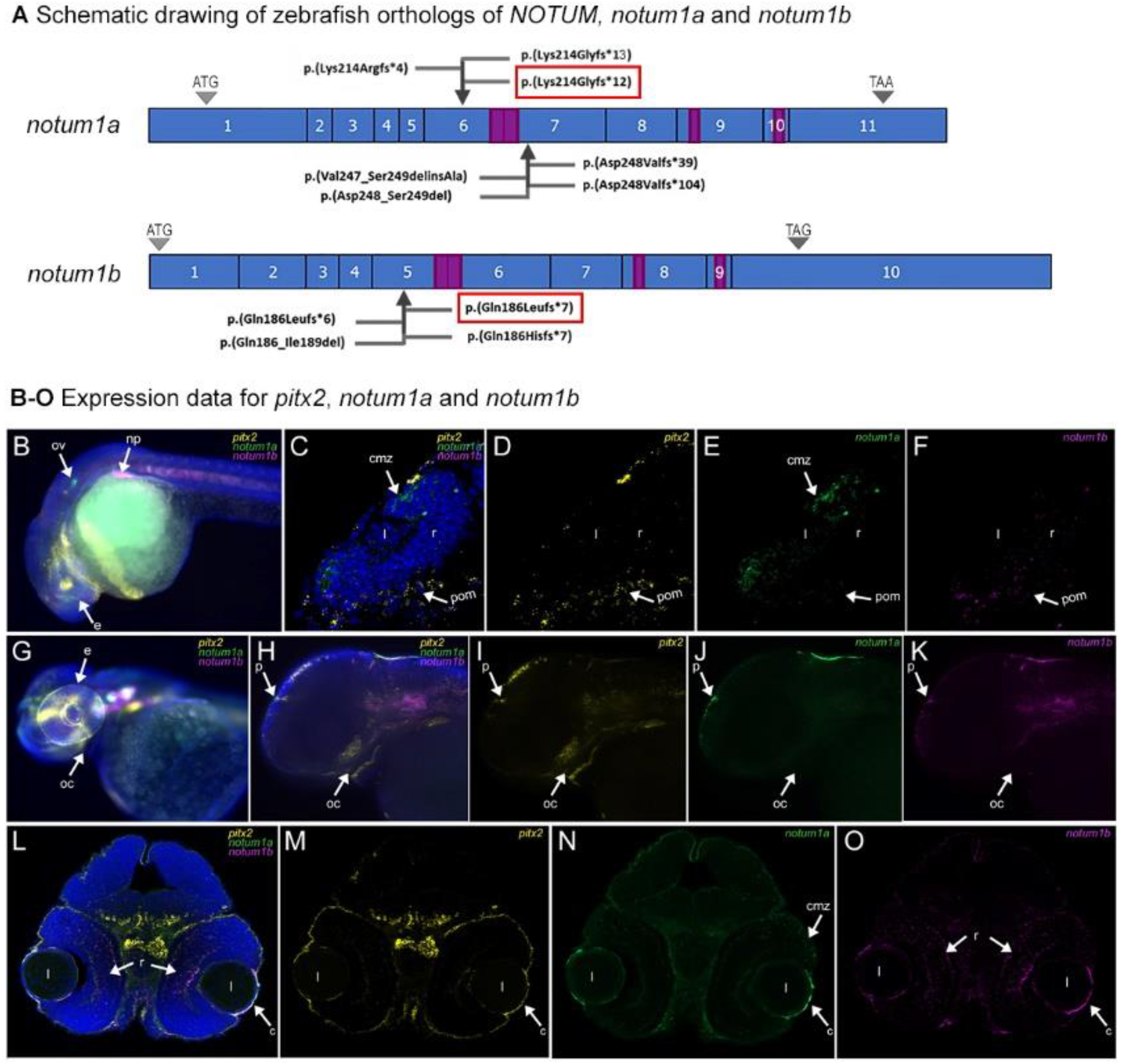
Schematics, CRISPR-mediated genomic editing, and expression patterns of *notum1a* and *notum1b.* **A)** Schematic drawing of zebrafish orthologs of *NOTUM, notum1a* and *notum1b.* CRISPR pairs were designed to target the regions encoding the active site motif and the serine of the catalytic triad of both *notum1a* and *notum1b* (first purple box). Localized genomic editing successfully occurred at three of the four target sites (arrows), resulting in multiple frameshift alleles in both genes. Red boxes denote the homozygous alleles present in *notum1*-deficient mutants (*notum1a^K214Gfs*12^; notum1b^Q186Lfs*7^).* **B-O)** Expression data for *pitx2* (yellow)*, notum1a* (green) and *notum1b* (magenta). RNAscope probes indicated broad expression patterns for both genes. Co-expression with *pitx2* was detected in the head mesenchyme and periocular region (especially for *notum1b*) at 24-hpf (B-F), as well as pharyngeal arches (*for notum1b*), pineal region, and developing cornea (for both *notum* transcripts) at 48-hpf (G-O). Divergent expression sites between *notum* homologs included the otic vesicle (enriched for *notum1a*) (B), nephric primordia (enriched for *notum1b*) (B), ciliary marginal zones (enriched for *notum1a* at both stages) (C, E, L, N) and central retina (enriched for *notum1b* at 48-hpf) (L, O). ov: otic vesicle; np: nephric primordia; e: eye; cmz: ciliary marginal zones; pom: periocular mesenchyme; l: lens; r: retina; oc: oral cavity (pharyngeal arches); p: pineal region; c: cornea.

Expression studies of *notum1a* and *notum1b* were performed using both digoxigenin-labeled anti-sense RNA probes and RNAScope probes (Figure 2B-O). Broad expression patterns, as illustrated by RNAscope, were observed for both genes. Co-expression with *pitx2* was detected in the head mesenchyme and periocular region (especially for *notum1b*) at 24-hpf (Figure 2B-F), as well as pharyngeal arches (*for notum1b*), pineal region, and developing cornea (for both *notum1* transcripts) at 48-hpf (Figure 2G-O). Divergent expression sites between *notum1* homologs included the otic vesicle (enriched for *notum1a*) (Figure 2B), nephric primordia (enriched for *notum1b*) (Figure 2B), ciliary marginal zones (enriched for *notum1a* at both stages) (Figure 2C, E, L, N) and central retina (enriched for *notum1b* at 48-hpf) (Figure 2L, O).

### Generation of notum1 single and double mutants

In order to determine the role of *notum1* during development, CRISPR pairs were designed to target the regions encoding the active site motif and the serine of the catalytic triad of both *notum1a* and *notum1b* (Figure 2A; Figure S1). Localized genomic editing successfully occurred at three of the four target sites. For *notum1a*, sequencing of offspring from mosaic fish resulted in five frameshift mutations: c.641delA, p.(Lys214Argfs*4); c.640_641delAA, p.(Lys214Glyfs*13); c.640_644delAAGGA, p.(Lys214Glyfs*12); c.743_746delinsTGTCTT, p.(Asp248Valfs*104); and c.743_749delinsTGCAAGACCTTTTGGAA, p.(Asp248Valfs*39). Two *notum1a* frameshift alleles also had additional mutations in cis but predicted to be downstream of protein truncation; c.743_748delACTCGG, p.(Asp248_Ser249del) was in cis with c.640_641delAA, p.(Lys214Glyfs*13), and c.740_745delTGGACT, p.(Val247_Ser249delinsAla) was in cis with c.640_644delAAGGA, p.(Lys214Glyfs*12). Sequence analysis of *notum1b* identified three frameshift mutations— c.556_557insTGATCTCTGATCATT, p.(Gln186Leufs*6); c.557_572delinsTCTTAACT, p.(Gln186Leufs*7); and c.558_565delGGAGGTCA, p.(Gln186Hisfs*7) — as well as one in-frame deletion— c.555_566delTCAGGAGGTCAT, p.(Gln186_Ile189del). The frameshift variants are predicted to disrupt the functional catalytic domains of both genes due to premature protein truncation. *notum1a* lines used for analysis were compound heterozygous for the c.743_749delinsTGCAAGACCTTTTGGAA, p.(Asp248Valfs*39) and c.640_641delAA, p.(Lys214Glyfs*13) alleles (*notum1a*^*D248Vfs*39*^/*notum1a*^*K214Gfs*13*^). *notum1b* lines were homozygous for the c.558_565delGGAGGTCA, p.(Gln186Hisfs^*7^) allele (*notum1b*^*Q186Hfs*7*^). *notum1* lines were homozygous for both the *notum1a* allele c.640_644delAAGGA, p.(Lys214Glyfs*12) and the *notum1b* allele c.557_572delinsTCTTAACT, p.(Gln186Leufs*7) (*notum1a*^*K214Gfs*12*^; *notum1b*^*Q186Lfs*7*^).

### Phenotypic characterization of notum1 single and double mutants

Single gene knockout lines for either *notum1a* (*notum1a*^*D248Vfs*39*^/*notum1a*^*K214Gfs*13*^) or *notum1b* (*notum1b*^*Q186Hfs*7*^) as well as a *notum1* double knockout line (*notum1a*^*K214Gfs*12*^; *notum1b*^*Q186Lfs*7*^) were examined for phenotypes. Consistent phenotypic presentation began at 5 dpf and manifested as an open jaw craniofacial deformity and an ectopic flared outgrowth in the gill region (Figure 3A-J’). In *notum1a*^*D248Vfs*39*^/*notum1a*^*K214Gfs*13*^ and *notum1b*^*Q186Hfs*7*^ mutants, both characteristics were variably expressed and incompletely penetrant, with *notum1a* single deficiency resulting in a more penetrant phenotype than *notum1b* single deficiency (25% jaw [17 of 68] and 84% gill [57 of 68] to 5% jaw [2 of 43] and 2% gill [1 of 43], respectively) (Figure 3B-C, G-H, G’-H’). By contrast, the phenotype was fully penetrant though still variably expressed in *notum1a*^*K214Gfs*12*^; notum1b^*Q186Lfs*7*^ double mutants (Figure 3D-E, I-J, I’-J’). Cartilage analysis by Alcian Blue staining displays anomalies primarily in the Meckel’s cartilage of the lower jaw (Figure 3F-J’). Alizarin Red staining of pharyngeal teeth performed in double mutants and wild-types revealed additional differences suggestive of a possible defect in tooth development (Figure 3K-N). All further analyses were focused on *notum1a*^*K214Gfs*12*^; *notum1b*^*Q186Lfs*7*^ double mutants hereinafter referred to as *notum1−/−*.

**Figure 3:**
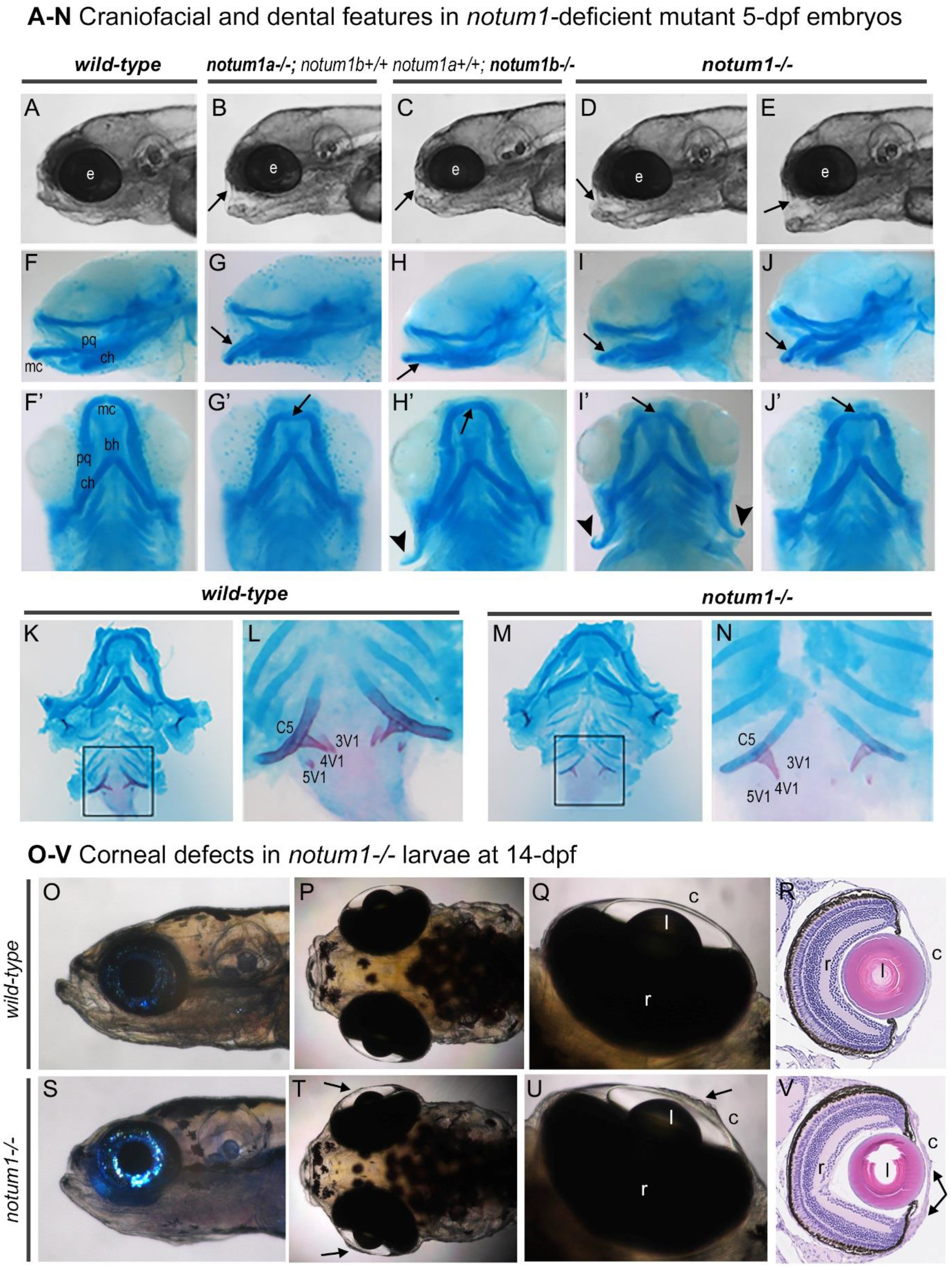
*notum1−/−* mutants display craniofacial, dental, and corneal defects at 5- and 14-dpf. **A-J’)** Craniofacial and dental features in *notum1−/−* mutant 5dpf embryos. In comparison to wild-type (A, F, F’), single gene knockout lines for either *notum1a* (B, G, G’) or *notum1b* (C, H, H’) as well as a *notum1* double knockout line (D-E, I-J, I’-J’) display an open jaw craniofacial abnormality (arrows in A-J) and an ectopic flared outgrowth in the gill region (arrowheads in F’-J’) beginning at 5 dpf. These features are variably expressed and incompletely penetrant in *notum1a* and *notum1b* single deficiency but fully penetrant in *notum1* double deficiency. Cartilage analysis by Alcian Blue staining displays anomalies primarily in the Meckel’s cartilage of the lower jaw (arrows in F-J’). **K-N)** Alizarin Red staining of pharyngeal teeth demonstrated a weaker staining pattern in 4 of 4 double mutants (M-N) vs. 1 of 4 wild-types (K-L), suggestive of a possible defect in tooth development. **O-V)** Corneal defects in *notum1−/−* mutant larvae at 14-dpf. At 14-dpf, *notum1−/−* mutants showed variable corneal anomalies, such as ballooning and unevenness (O-Q, arrows in S-U). Sectioning demonstrated defects in corneal thickness, ranging from extremely thin, most notably at the periphery, to unusually thick (R, arrows in V). e: eye; mc: Meckel’s cartilage; pq: palatoquadrate; ch: ceratohyal; bh: basihyal; c5: ceratobranchial 5; 3V1-5V1: pharyngeal teeth; l: lens; r: retina; c: cornea.

No consistent differences in ocular structures were detected between wild-type and *notum1−/−* mutants by gross observation, imaging or histology at 5-dpf. Analysis at later timepoints revealed the onset of an ocular phenotype. At 14-dpf, *notum1−/−* mutants showed variable corneal anomalies, such as ballooning and unevenness (Figure 3O-Q, S-U). Sectioning demonstrated defects in corneal thickness, ranging from extremely thin, most notably at the periphery, to unusually thick (Figure 3R, V). Examination of *notum1−/−* adult fish revealed the persistence of ocular defects. At 7-mpf and 14-mpf, no obvious differences were observed by light microscopy between wild-types and *notum1−/−* mutants. Histology and subsequent staining, however, revealed some striking features (Figure 4A-L). Regarding ocular structures, a spectrum of corneal disruption is evident. The corneal epithelium is of uneven thickness, even to the point of absence in portions of some corneas; in addition, the corneal stroma appears loose and disorganized with fibrous or vacuous inclusions appearing between the stromal layers, particularly at the junction with the annular ligament (Figure 4A-L). Periodic Acid Schiff staining (PAS) also suggests that mucin-secreting goblet cells may be present in some eyes beyond the annular ligament and into the cornea (data not shown).

**Figure 4:**
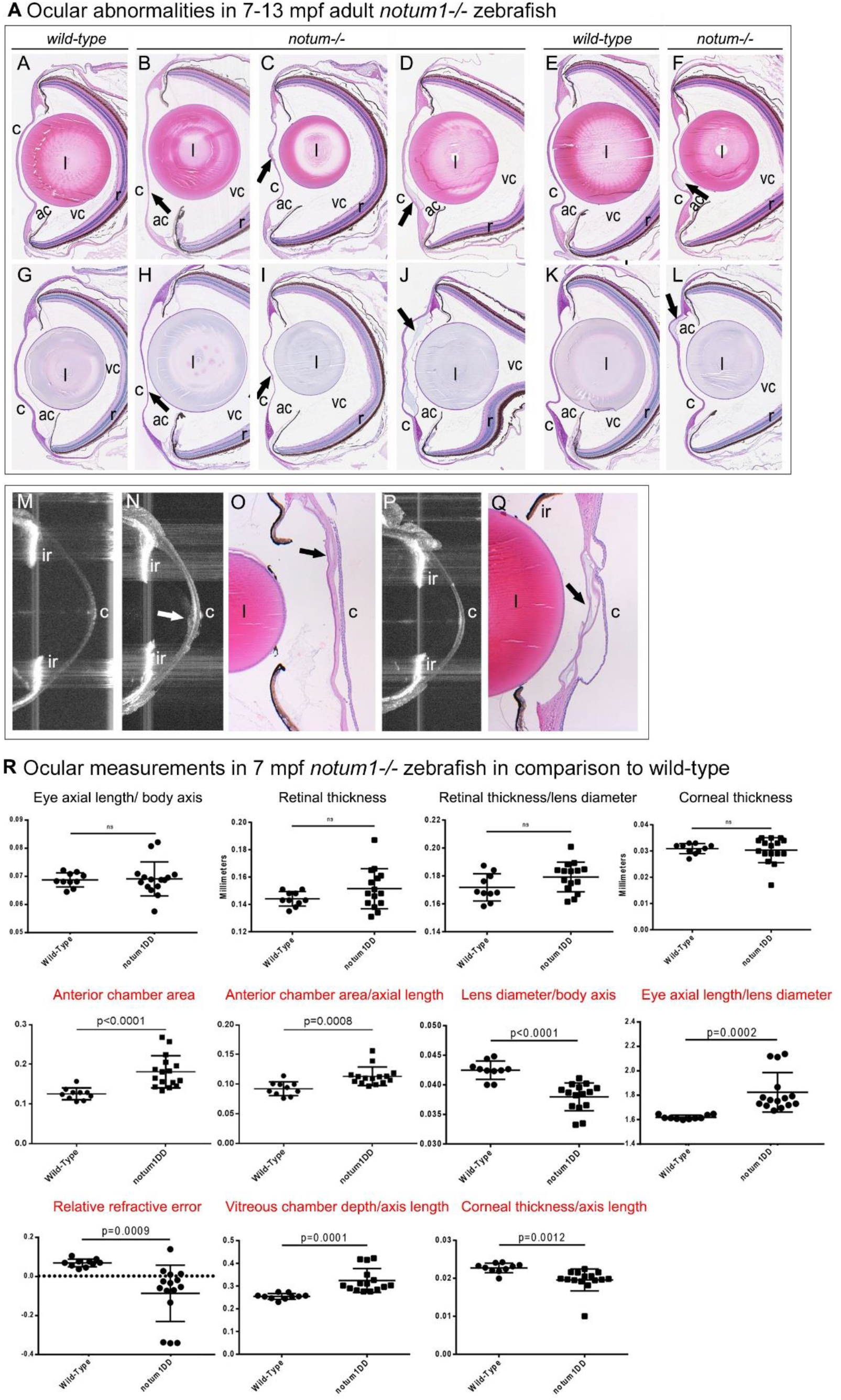
Ocular anomalies and measurements of adult *notum1−/−* mutants. **A-L)** Ocular abnormalities in 7-mpf (B-D, H-J) and 13-mpf (F, L) *notum1−/−* mutants as compared to wild-type 7-mpf (A, G) and 13-mpf (E, K) counterparts. The spectrum of corneal disruption involves the corneal epithelium being of uneven thickness, even to the point of absence in portions of some corneas; in addition, the corneal stroma appears loose and disorganized with fibrous or vacuous inclusions appearing between the stromal layers, particularly at the junction with the annular ligament (arrows in A-L). **M-Q)** In vivo analysis of 7 and 8-mpf wild-type (M) and *notum1−/−* (N-Q) adult eyes by OCT. OCT B-scans demonstrated fibrotic deposits and other abnormalities occasionally seem in *notum1−/−* corneas (arrow in M-N, P). Anomalies detected by OCT were confirmed by sectioning of the same eyes (arrows in O, Q). **R)** Ocular measurements in 7-mpf *notum−/−* zebrafish in comparison to wild-type. *notum1−/−* mutant eyes displayed greater anterior chamber areas when normalized to the axial length of the eye and by direct area comparison. Vitreous chamber depth was also greater in . *notum1−/−* mutants when normalized to body axis length. Lenses of *notum1−/−* mutant eyes were smaller than wild-types when normalized to body axis length, while axial lengths did not change. *notum1−/−* mutants were also myopic, as indicated by differences in relative refractive error. Finally, *notum1−/−* mutant corneas were thinner when normalized to the axial length of each eye. F-tests for variance associated with the statistical analyses (unpaired t-tests with Welch’s correction) showed that the following *notum 1−/−* eye metrics displayed greater variance than wild-types: normalized axial length (normalized to either body axis or lens diameter), relative refractive error, vitreous chamber depth (normalized to either body axis or lens diameter), retinal thickness, corneal thickness, and anterior chamber area. l: lens; ac: anterior chamber; vc: vitreous chamber; r: retina; c: cornea; ir: iris.

In vivo analysis of 7 and 8-mpf wild-type and *notum1−/−* adult eyes was conducted using OCT (Figure 4M-R). Selected movies were generated from adjacent OCT B-scans to demonstrate fibrotic deposits and other abnormalities seem in *notum1−/−* corneas (Supplemental Movies 1 and 2). Anomalies detected by OCT were confirmed by sectioning of the same eyes (Figure 4 M-Q). Measurements were taken using OCT images and used to determine various ocular parameters in mutant eyes in comparison to wild-type. The *notum1−/−* mutant eyes displayed greater anterior chamber areas relative to their wild-type counterparts when normalized to the axial length of the eye and by direct area comparison (Figure 4R). Vitreous chamber depth was also greater in *notum1−/−* mutants when normalized to body axis length (Figure 4R). Lenses of *notum1−/−* mutant eyes were smaller than wild-types when normalized to body axis length, while axial lengths did not change (Figure 4R). *notum1−/−* mutants were also myopic, as indicated by differences in relative refractive error (Figure 4R) (Collery et al., 2014). Finally, *notum1−/−* mutant corneas were thinner when normalized to the axial length of each eye (Figure 4R). F-tests for variance associated with the statistical analyses (unpaired t-tests with Welch’s correction) showed that the following *notum 1−/−* eye metrics showed greater variance than wild-types: normalized axial length (normalized to either body axis or lens diameter), relative refractive error, vitreous chamber depth (normalized to either body axis or lens diameter), retinal thickness, corneal thickness, and anterior chamber area (Figure 4R).

Immunostaining for corneal keratin sulfate proteoglycan (CKS) (a marker for corneal stroma) (Zhao et al., 2006) and n-cadherin (cdh-2) (stains the corneal endothelium and epithelium as well as and lens epithelial layer) (Erdmann, Kirsch, Rathjen, & More, 2003; Lu et al., 1999) further illustrated the epithelial and stromal anomalies in *notum1−/−* mutants (Figure 5A-I). At 14-dpf, staining demonstrated notable corneal thinning and near loss of the corneal epithelium in some eyes (Figure 5B); in addition, aberrant CKS staining appeared in the corneal epithelium of some eyes, suggestive of compromised epithelial identity (Figure 5C-D). At 7- and 13-mpf, *notum1−/−* mutants continue to display corneal epithelial unevenness as well as disorganized, fibrous stromal anomalies, thus providing additional support for the importance of *notum1* in the development and maintenance of proper corneal identity (Figure 5E-I).

**Figure 5:**
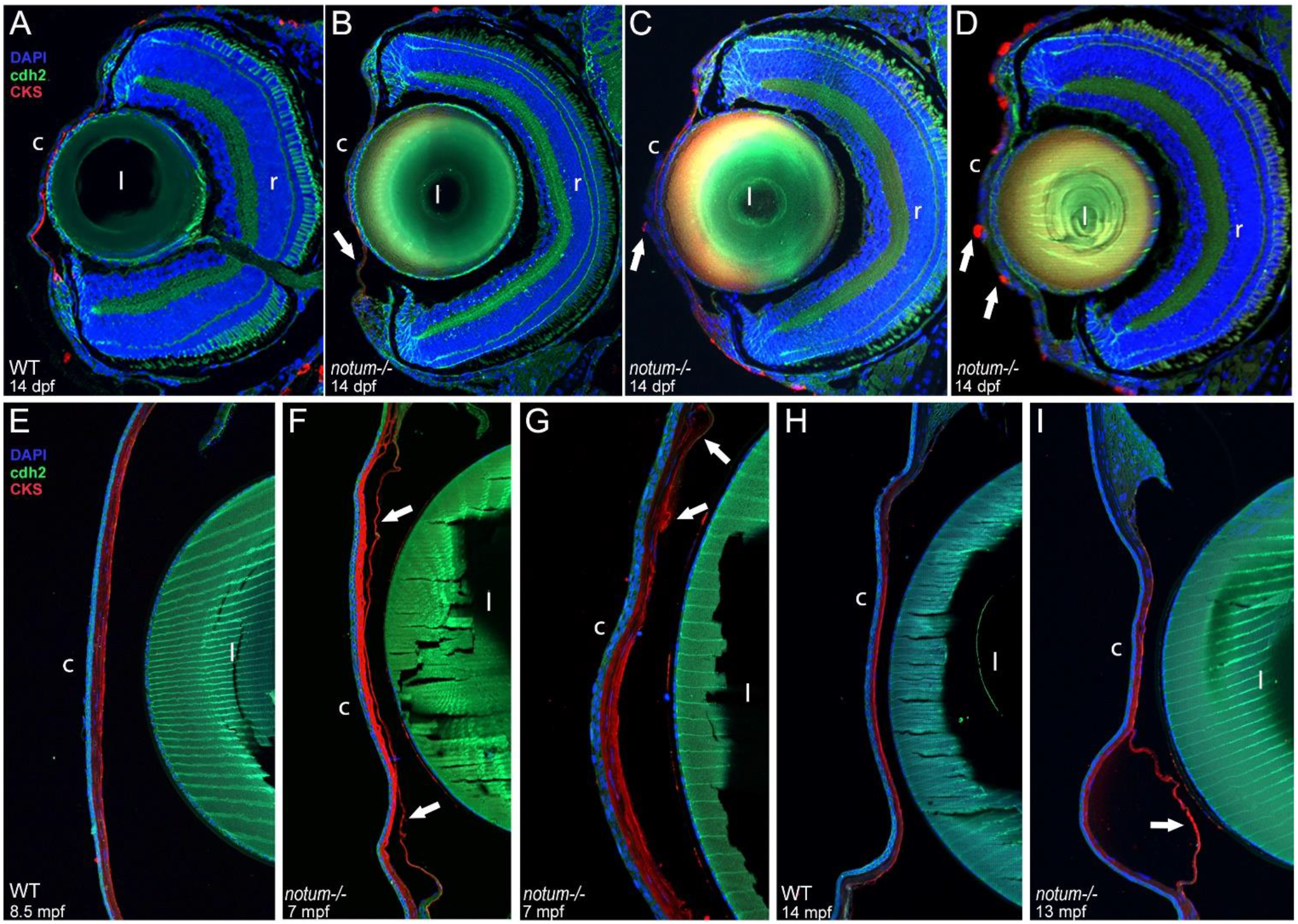
Immunohistochemical analysis of the *notum1−/−* mutant ocular phenotype. **A-D)** Immunostaining for the corneal stroma marker CKS (red), the corneal endothelium and cornea and lens epithelial marker cdh-2 (green), and DAPI (blue) in 14-dpf wild-type (A) and *notum1−/−* mutant (B-D) corneas. Staining demonstrated notable corneal thinning and near loss of the corneal epithelium in some eyes (arrow in B); in addition, aberrant CKS staining appeared in the corneal epithelium of some eyes, suggestive of compromised epithelial identity (arrows in C-D). **E-I)** CKS, cdh-2, and DAPI staining of wild-type corneas at 8.5-mpf (E) and 14-mpf (H) and *notum1 −/−* mutant corneas at 7-mpf (F-G) and 13-mpf (I). *notum1−/−* mutants continue to display corneal epithelial unevenness as well as disorganized, fibrous stromal anomalies (arrows in F-G, I). c: cornea; l: lens; r: retina.

To delve into the ultrastructural features underlying the corneal phenotypes, electron microscopy was performed on 8-mpf wild-type and *notum1−/−* adult eyes (Figure 6A-F). Of particular interest was the condition of the corneal endothelium, as this structure is difficult to visualize with histology. *notum1−/−* mutant corneas did possess an endothelial layer; however, this layer demonstrated multiple abnormalities. For instance, the junction of the endothelium with the annular ligament was disrupted in the *notum1−/−* mutant (Figure 6A, C). In the wild-type iridocorneal angle, the endothelial layer gradually begins to stratify into multiple organized layers before joining the annular ligament (Figure 6A). In the mutant angle, the stratifying layers separate from one another and become highly disorganized before reaching the annular ligament (Figure 6C). The wild-type central corneal endothelium appears as a monolayer of uniform thickness; in the *notum1−/−* mutant, the central corneal endothelial layer fluctuates in thickness and appears abnormal, unhealthy, and vacuous (Figure 6B, D). Finally, the endothelial layer covering the annular ligament takes on an unusual, compact morphology compared to the wild-type (Figure 6E-F). Therefore, the ultrastructural data reveals anomalies in the corneal endothelium, thus confirming abnormalities in all three corneal layers in *notum1−/−* mutants.

**Figure 6:**
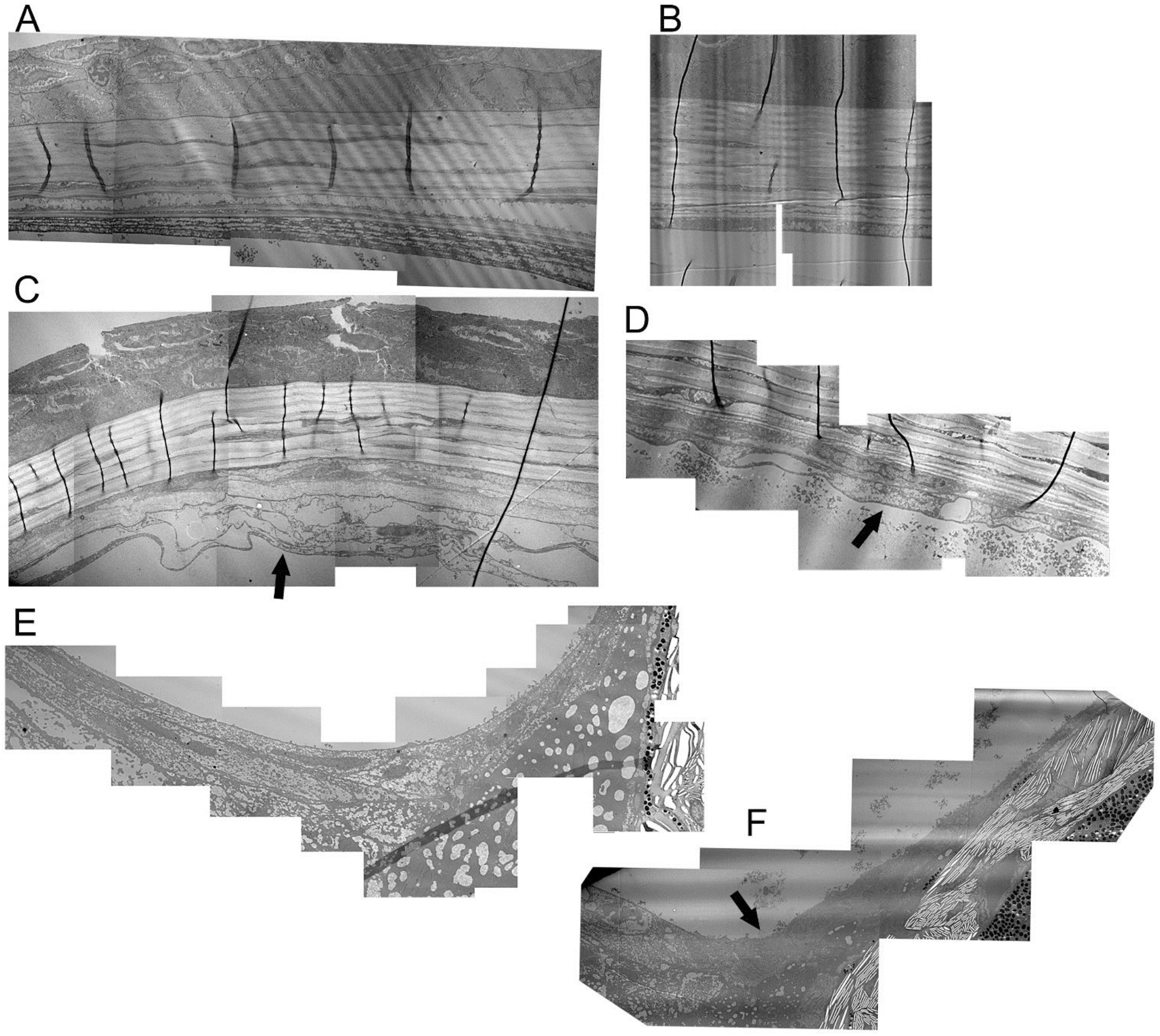
Electron microscopy of *notum1−/−* mutant adult anterior segments. **A-F)** 8-mpf wild-type (A-B, E) and *notum1−/−* mutant (C-D, F) iridocorneal angles (A, C, E-F) and central corneas (B, D). The junction of the endothelium with the annular ligament was disrupted in the *notum1−/−* mutant (A, C). In the wild-type iridocorneal angle, the endothelial layer gradually begins to stratify into multiple organized layers before joining the annular ligament (A); in the mutant angle, the stratifying layers separate from one another and become highly disorganized before reaching the annular ligament (arrow in C). The wild-type central corneal endothelium appears as a monolayer of uniform thickness (B); in the *notum1−/−* mutant, the central corneal endothelial layer fluctuates in thickness and appears abnormal, unhealthy, and vacuous (arrow in D). The endothelial layer covering the annular ligament also takes on an unusual, compact morphology compared to the wild-type (arrow in E-F).

### Pathways affected by notum1 deficiency

To examine the molecular effects of *notum1* deficiency, microarray analysis was performed on 24-hpf wild-type and *notum1−/−* whole heads. 1170 down-regulated and 860 up-regulated targets were found to be significantly differentially expressed. Combined annotation via Biomart, cross referencing of previous arrays, and IPA recognized 508 down-regulated and 393 up-regulated transcripts as having human orthologs. Signaling pathways predicted to display increased activity include WNT/β-catenin, as would be anticipated when a WNT antagonist like *NOTUM* is disrupted, Hippo, cardiac β-adrenergic, ERK/MAPK, and PI3K/AKT signaling (Tables S3 and S4). Pathways predicted to exhibit decreased activity include calcium, integrin, and androgen signaling as well as inhibition of matrix metalloproteinases (Table S3). Increased activity was predicted in broader processes including connective tissue tumors, infarction, quantity of blood cells, release of catecholamine, and organismal morbidity or mortality (Table S3). Processes such as cell and neurite branching, neuronal and synapse development, formation of intercellular and gap junctions, and cytoplasm and cytoskeleton organization all displayed indications of decreased activity (Table S3). Full microarray data will be available pending submission to the ArrayExpress database at EMBL-EBI (www.ebi.ac.uk/arrayexpress).

## DISCUSSION

Studies into downstream pathways of PITX2/pitx2 identified *notum1* (*notum1a* and *notum1b*) as significantly affected transcripts. Notum is a secreted carboxylesterase that is a member of the α/β hydrolase protein family and shares high conservation around its active site motif (G-X-S-X-G) and the family-characteristic Ser, Asp, His catalytic triad (Flowers et al., 2012a; Giraldez, Copley, & Cohen, 2002; Holmquist, 2000; Kakugawa et al., 2015). When Notum was discovered, it was believed to be a glypican-specific phospholipase that worked to inhibit signaling by cleaving the glycosylphosphatidylinositol (GPI) anchor of glypicans (Kreuger, Perez, Giraldez, & Cohen, 2004; Traister, Shi, & Filmus, 2008). In 2015 it was shown that Notum interacts with glypicans through sulfated glycosaminoglycan chains in order to perform its primary function of specifically cleaving the O-palmitoleate moiety of Wnt proteins (Kakugawa et al., 2015; Zhang et al., 2015). Thus, Notum was the first described extracellular protein deacylase. Interestingly, though the Wnt pathway has been the primary focus of study, the hydrophobic pocket of NOTUM was shown to have a preference for multiple substrates beyond the Wnt-associated *cis*-unsaturated myristoleic (14 carbons) and palmitoleic (16 carbons) acids, namely saturated 8- to 12-carbon linear carboxylic acids; in addition, the carboxyester bond to be cleaved can either be a carboxy-oxoester or a carboxy-thioester bond (Kakugawa et al., 2015). This raises the possibility that NOTUM may also act on other yet unidentified targets in addition to its clear preference for Wnt proteins.

Notum has previously been studied in other animal models including zebrafish, where *notum1a* knockdown and over-expression demonstrated *notum1a’s* importance in head induction within the first day of development (Flowers et al., 2012a). Our data from genetic *notum1a−/−* or even complete *notum1−/−* mutants show a significantly less severe phenotype; similar discrepancies between morpholino and genetic mutant data has been discussed in many papers and may be due to several factors (Kok et al., 2015; Law & Sargent, 2014; Rossi et al., 2015; Schulte-Merker & Stainier, 2014; Stainier, Kontarakis, & Rossi, 2015) . The initial discovery of *Notum* occurred in Drosophila, where complete loss of notum was found to be lethal during pupal stages (Gerlitz & Basler, 2002). Notum was found to play a key role in wing development and maintaining the Drosophila Wingless (mammalian WNT1) signaling gradient (Gerlitz & Basler, 2002; Giraldez et al., 2002). Notum was also shown to modulate synapse growth, architecture, ultrastructural development, and functionality in flies (Kopke, Lima, Alexandre, & Broadie, 2017). *Notum*-deficient mouse lines have been generated and exhibited differing phenotypes. One line found *Notum−/−* mice to display dentin dysplasia and reduced viability, with one quarter also displaying kidney agenesis (Vogel et al., 2016). Another line confirmed *Notum* homozygous perinatal lethality due to abnormal kidney development while also demonstrating that Notum was essential for proper patterning of tracheal mesenchyme (Gerhardt et al., 2018).

Our *notum1−/−* line represents the first zebrafish model to knockout both *notum1* homologs and examine the long-term effects of *notum1* deficiency. *notum1−/−* mutants display craniofacial defects including a possible delay in tooth development and an unusual gill structure anomaly; these features are reminiscent of some aspects of mouse *Notum* deficiency, namely dentin dysplasia and perhaps tracheal patterning. Ocular features have a later onset and include corneal anomalies in all three layers and disproportionally small lens resulting in myopia. These defects overlap features associated with *pitx2/Pitx2* deficiency including loss of corneal integrity, jaw and dental anomalies (Gage et al., 1999; Hendee et al., 2018; Ji, Buel, & Amack, 2016; Li et al., 2014) as well as *Dkk2* deficiency, which also displays compromised corneal identity (Gage et al., 2008; Mukhopadhyay et al., 2006). The presented data suggest that PITX2/pitx2 acts on multiple targets (*NOTUM/notum1, DKK2/dkk2*) to negatively affect the WNT pathway during anterior segment development and their combined function is required for the proper eye formation.

Pathway analysis of differentially expressed targets in *notum1* mutants suggests an overall increase in WNT/β-catenin signaling. Wnt signaling is a key mechanism of directing cell proliferation, cell polarity and cell fate determination during embryonic development (MacDonald, Tamai, & He, 2009). In canonical Wnt signaling, the absence of Wnt ligands results in the sequestration of β-catenin in the cytoplasm as part of a destruction complex with Axin, APC, GSK3, CK1 and PP2A; β-catenin is phosphorylated by CK1 and GSK3 and thereby targeted for ubiquitination and proteasomal degradation. In the presence of Wnt ligands, a receptor complex forms between Frizzled and LRP5/6. Disheveled recruitment by Frizzled leads to LRP5/6 phosphorylation and Axin recruitment, which disrupts the Axin-mediated phosphorylation and degradation of β-catenin and allows β-catenin to accumulate in the nucleus where it serves as a co-activator for TCF to activate Wnt responsive genes (MacDonald et al., 2009).

*notum1* mutants display down-regulation of multiple players in this pathway that serve to modulate both enhancement and repression of Wnt signaling. Down-regulation of Frizzled receptor 5 (FZD5) would reduce substrates available for WNT ligand binding. Casein kinase 2 alpha 1 (CSNK2A1) down-regulation could reduce phosphorylation and activation of the Disheveled complex, which would reduce inhibition of the destruction complex and increase phosphorylation and sequestration of β-catenin in the cytoplasm. Four phosphatases—protein phosphatase, Mg2+/Mn2+ dependent IL (PPM1L) and protein phosphatase 2 regulatory subunits Bbeta (PPP2R2B), Bgamma (PPP2R2C) and B’epsilon (PPP2R5E)—were also downregulated and involved in several processes, namely the protein phosphatase 2a (PP2A) complex that participates in the β-catenin destruction complex. B56 (B’) subunits [as in PPP2R5E] increase the phosphorylation-induced proteasomal degradation of β-catenin (Seeling et al., 1999); B subunits are not indicated to act through the exact same mechanism as B’ subunits but still regulate phosphatase specificity and may behave in a similar manner. IPA predictions indicated that the affected phosphatases would be down-regulated when the Wnt pathway is active, so their down-regulation in *notum1* mutants corresponds with higher Wnt activity. Selective down-regulation of Sox8 could reduce inhibition of β-catenin/LEF/TCF transcriptional activity. SRC proto-oncogene, non-receptor tyrosine kinase (SRC) is significantly down-regulated and also a mediator of cadherin/β-catenin interaction. Phosphorylation of cadherin and β-catenin by SRC family members results in dissociation of the cadherin/β-catenin complex and increased β-catenin in the cytoplasm available for signaling (Behrens et al., 1993; Qi, Wang, Romanyuk, & Siu, 2006). The reduction in SRC activity would likely stabilize the cadherin/β-catenin complex and reduce the pool of free β-catenin. Thus, the predicted net result of an increase in Wnt signaling, which is consistent with loss of a Wnt antagonist, may be mediated by down-regulation of lesser known Wnt pathway members in *notum1*-deficient mutants.

In addition, the cadherin/β-catenin complex helps to promote cell adhesion. In the case of epithelial cells, E-cadherin and β-catenin interact to link the cell membrane to microtubules within the cell and to form junctions with neighboring cells (Tian et al., 2011). Casein kinase 2 phosphorylation of E-cadherin and β-catenin stabilizes the complex and preserves cell adhesion, while phosphorylation by SRC destabilizes the complex and disrupts cell adhesion (Behrens et al., 1993; Bek & Kemler, 2002; Lickert, Bauer, Kemler, & Stappert, 2000; Tian et al., 2011). Another factor that may come into play are matrix metalloproteinases (MMPs). MMPs have been implicated in disrupting epithelial cell integrity (Tan et al., 2010; Tian et al., 2011; Zheng et al., 2009). The results of the microarray indicate that five MMPs are significantly upregulated in the *notum1* mutant heads. Therefore, it is possible that the observed phenotype of compromised corneal epithelial integrity in *notum1*-deficiency could be due in part to the destabilization of E-cadherin/β-catenin interaction resulting from a combination of down-regulation of CSNK2A1 and up-regulation of several MMPs overwhelming a predicted stabilization generated by down-regulation of SRC.

In summary, in *notum1* we have identified and characterized a novel downstream member of the pitx2 pathway essential for proper embryonic development. *NOTUM/notum1* may serve as a new causative gene for unexplained human ocular and/or craniofacial disorders.

## ACKNOWLEDGEMENTS

We would like to thank Dr. Eric Weh for CRISPR design guidance and injection aid and Rebecca Tyler for sequencing assistance. We would also like to thank the Department of Cell Biology, Neurobiology & Anatomy for use of shared equipment, as well as Christine Duris at the Children’s Research Institute Histology Core and Clive Wells and Robert Goodwin at the Medical College of Wisconsin Electron Microscopy Facility for technical assistance.

## FUNDING

This work is supported by awards from National Institutes of Health R01EY015518 (EVS), and T32EY014537 (KEH), and Children’s Research Institute Foundation at Children’s Hospital of Wisconsin (EVS).

## CONFLICT OF INTEREST

The authors have no conflict of interest to disclose.

