## Supplemental Materials for "*notum1*, acting downstream of pitx2, is essential for proper eye and craniofacial development"

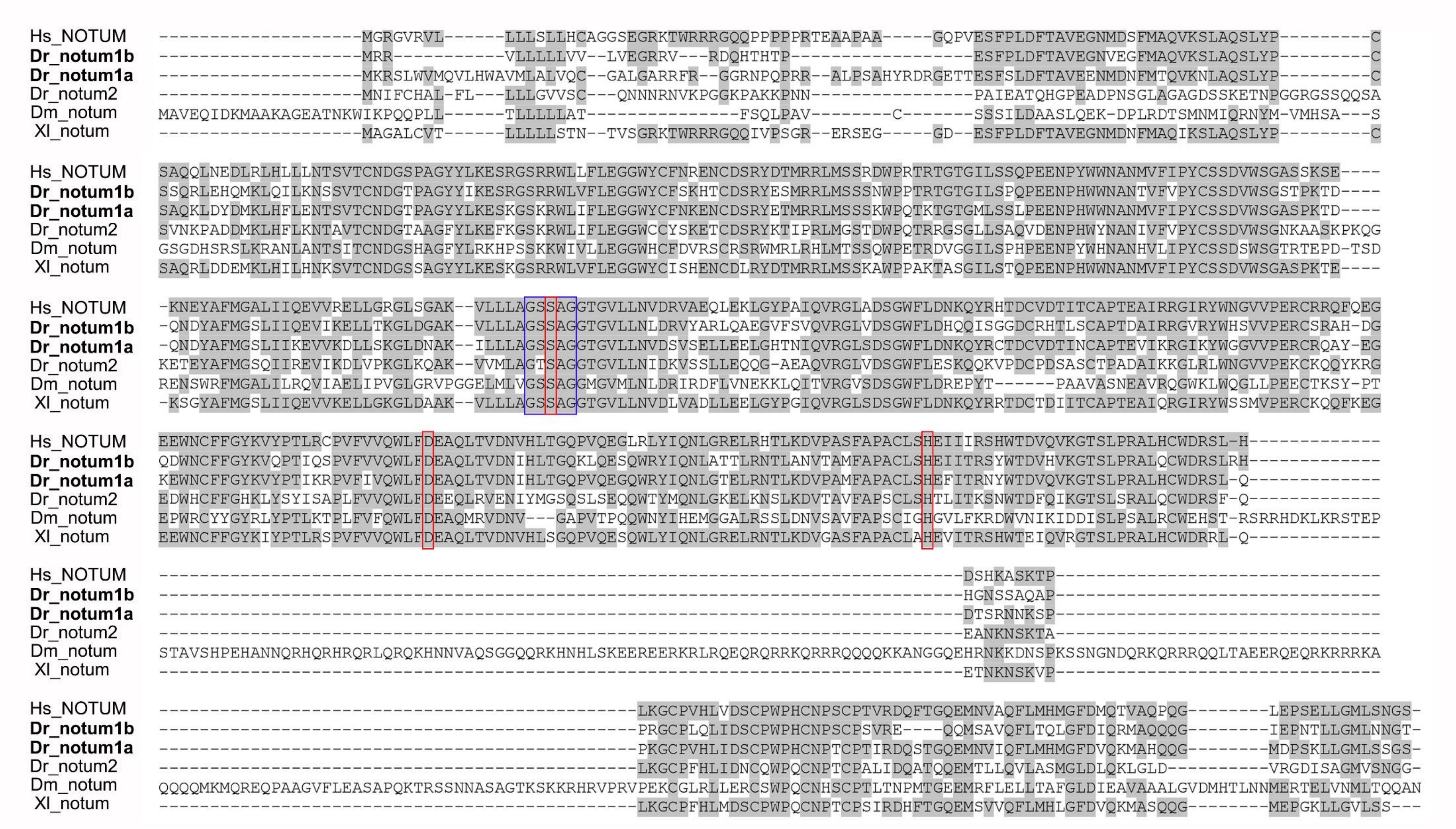


**Figure S1: Protein alignments of NOTUM homologs.** Amino acid alignments of: human (Homo sapien) NOTUM; zebrafish (Danio rerio) notum1b, notum1a, and notum2; fruit fly (Drosophila melanogaster) notum; and African clawed frog (Xenopus laevis) notum. Note the high conservation around the active site motif G-X-S-X-G (blue box) and the Ser, Asp, His catalytic triad (red boxes) characteristic of the α/β hydrolase protein family.

**Table S1: Oligonucleotides utilized for qRT-PCR, clone sequencing, ISH, sgRNA synthesis, and genotyping.**

| Gene | Sequence (5'→3') | Product size |
|---|---|---|
| ***Zebrafish primers for quantitative reverse transcription PCR*** | | |
| *notum1a* | CACTGACTGTGTGGACACCA (forward) | 109 bp |
| *notum1a* | TCCTTCATAAGCCTGCCTGC (reverse) | |
| *notum1b* | CCAACGTCACGGCCATGTTT (forward) | 192 bp |
| *notum1b* | TGTCAATCAACTGCAGCGGG (reverse) | |
| *dkk2* | GCAGCAACTACATCTGCATTCC (forward) | 150 bp |
| *dkk2* | CTTCGTGACCTTTGAGAGAGATC (reverse) | |
| *β-actin* | GAGAAGATCTGGCATCACAC (forward) | 324 bp |
| *β-actin* | ATCAGGTAGTCTGTCAGGTC (reverse) | |
| | ***Zebrafish primers for clone sequencing and in situ probe generation*** | |
| *notum1a* | GGTGATGCTGGCTTTGGTTC (forward) | 623 bp |
| *notum1a* | AGAGCAGATCCTTCACGACC (reverse) | |
| *notum1b* | GCGGCTCTACACCAAAGACT (forward) | 756 bp |
| *notum1b* | TGTCAATCAACTGCAGCGGG (reverse) | |
| *notum1b* | AGGAGCTGCTGTGTGAGATG (forward 1) | 1424 bp [full cDNA] |
| *notum1b* | TCTAGGTGCCGTTATTGAGC (reverse 1) | |
| *notum1b* | TGAACCTGGACCGTGTGTAT (forward 2) | 763 bp [cDNA 3' end] (with reverse 1) |
| *notum1b* | CTGATCTGCTGATGATCCAG (reverse 2) | 770 bp [cDNA 5' end] (with forward 1) |
| ***Zebrafish primers for CRISPR sgRNA generation*** | | |
| *notum1a_ex6* | TAGGGTTCACTGATCATAAAGG (forward) | |
| *notum1a_ex6* | AAACCCTTTATGATCAGTGAAC (reverse) | |
| *notum1a_ex7* | TAGGTCTTGCTGAATGTGGACT (forward) | |
| *notum1a_ex7* | AAACAGTCCACATTCAGCAAGA (reverse) | |
| *notum1b_ex5* | TAGGGCTCTCTGATCATTCAGG (forward) | |
| *notum1b_ex5* | AAACCCTGAATGATCAGAGAGC (reverse) | |
| *notum1b_ex8* | TAGGTTCAGTGGCTGTTTGACG (forward) | |
| *notum1b_ex8* | AAACCGTCAAACAGCCACTGAA (reverse) | |
| ***Zebrafish primers for line genotyping*** | | |
| *notum1a_ex6* | GATCCCCAGTAACATGAGTTC (forward) | 404 bp |
| *notum1a_ex6* | GCAGAATGCACACACAAGCT (reverse) | |
| *notum1a_ex7* | ATGTAAGTATGTGCTCCTCAGTG (forward) | 215 bp |
| *notum1a_ex7* | ACTTGATTCCTCTCTTGATGACC (reverse) | |
| *notum1b_ex5* | CGGCTCTACACCAAAGACTG (forward) | 218 bp |
| *notum1b_ex5* | TCTCTCACCTGCTTCCGG (reverse) | |
| *notum1b_ex8* | TCTTCGGCTACAAGGTCCAG (forward) | 340 bp |
| *notum1b_ex8* | TTGGATCCTGCTTGTGTCTG (reverse) | |
| *notum2* | TCCGGAAAAGTGTAAGCAGC (forward) | 304 bp |
| *notum2* | TCCCCAGGTTCTGCATGTAG (reverse) | |

**Table S2: Signaling pathways and developmental/cellular processes predicted to be affected in (*CE4:GFP):pitx2*$^{M64*}$ vs. (*CE4:GFP):WT* GFP-positive cell populations.**

| IPA prediction | Increased activity | Decreased activity |
|---|---|---|
| **Signaling pathways** | Gp6 | Adrenomedullin<br>Protein Kinase A (PKA) |
| **Developmental and cellular processes (subset)** | Dopamine concentration<br>Brain growth<br>Head development<br>Sensory organ development<br>Abdomen development | Cellular attachment<br>Kidney development |

**Table S3: Signaling pathways and developmental/cellular processes predicted to be affected in wild-type vs. *notum1-/-* embryos.**

| IPA prediction | Increased activity | Decreased activity |
|---|---|---|
| **Signaling pathways (subset)** | WNT/β-catenin<br>Hippo<br>Cardiac β-adrenergic<br>ERK/MAPK<br>PI3K/AKT | Calcium<br>Integrin<br>Androgen<br>Inhibition of matrix metalloproteinases |
| **Developmental and cellular processes (subset)** | Connective tissue tumors<br>Infarction<br>Quantity of blood cells<br>Release of catecholamine<br>Organismal morbidity or mortality | Cell branching<br>Neurite branching<br>Neuronal development<br>Synapse development<br>Intercellular junction formation<br>Gap junction formation<br>Cytoplasm organization<br>Cytoskeleton organization |

**Table S4: Transcripts identified by IPA as differentially regulated in *notum-/-* head tissues (part of WNT pathway).**

| Gene | Entrez Gene Name | Ensembl | Expr Log Ratio | Expected | Type(s) |
|---|---|---|---|---|---|
| *PPM1L* | protein phosphatase, Mg2+Mn2+ dependent 1L | ENST00000497343 | -2.613 | Down | phosphatase |
| *PPP2R2B* | protein phosphatase 2 regulatory subunit Bbeta | ENSG00000156475 | -1.012 | Down | phosphatase |
| *PPP2R2C* | protein phosphatase 2 regulatory subunit Bgamma | ENSG00000074211 | -1.052 | Down | phosphatase |
| *PPP2R5E* | protein phosphatase 2 regulatory subunit B'epsilon | ENSG00000154001 | -1.016 | Down | phosphatase |
| *SOX8* | SRY-box 8 | ENSG00000005513 | -1.141 | Down | transcription regulator |
| *SRC* | SRC proto-oncogene, non-receptor tyrosine kinase | ENSG00000197122 | -2.157 | Down | kinase |
| *CSNK2A1* | casein kinase 2 alpha 1 | ENSG00000101266 | -1.592 | Up | kinase |
| *FZD5* | frizzled class receptor 5 | ENSG00000163251 | -1.605 | Up | G-protein coupled receptor |
| *MDM2* | MDM2 proto-oncogene | ENSG00000135679 | 1.482 | | transcription regulator |
| *NR5A2* | nuclear receptor subfamily 5 group A member 2 | ENSG00000116833 | -1.627 | Up | ligand-dependent nuclear receptor |
